# PhytoFam: A Nextflow Pipeline for Genome-Wide Analysis of Plant Gene Families

**DOI:** 10.64898/2026.07.23.740317

**Authors:** Sanam Parajuli, Bibek Adhikari, Anne Fennell, Madhav P. Nepal

**Author notes:** Corresponding author: Madhav P Nepal, ^1^Department of Biology and Microbiology, South Dakota State University, Brookings, SD.

## Abstract

Genome-wide identification of plant gene families is essential for functional and evolutionary studies but often requires the use of multiple independent tools for homolog detection, domain validation, orthology assignment, and phylogenetic analysis. This fragmented approach involves extensive manual scripting, complicates reproducibility and parameter tracking, and may require additional steps to remove redundant protein isoforms. To address these challenges, we developed PhytoFam, a Nextflow-based workflow that automates gene family identification from proteome input through phylogenetic reconstruction. The pipeline integrates HMMER for candidate sequence identification, isoform-aware deduplication, InterProScan for domain confirmation, BLAST reciprocal best hit (RBH) analysis for orthology assignment, MUSCLE for multiple sequence alignment with optional outgroup incorporation, TrimAl for alignment trimming, and IQ-TREE3 for phylogenetic reconstruction. PhytoFam is portable across local workstations and high-performance computing environments and supports deployment through Conda, Docker, and Singularity. We validated the workflow using the *Morus alba* MADS-box gene family, where the complete analysis finished in 1 h 10 min (9 CPU h). IQ-TREE3 accounted for most of the execution time, whereas InterProScan showed the highest memory requirement with a peak resident set size of 4.5 GB. PhytoFam provides a reproducible, automated, and scalable solution for plant gene family identification and phylogenetic analysis. The pipeline is freely available at https://github.com/sanamparajuli/PhytoFam.

## 1 Introduction

Plant genomes exhibit extensive gene family expansion, driven by polyploidy events, tandem duplication, and transposon-mediated rearrangements (Panchy et al. 2016; Wendel et al. 2016). These gene families underpin natural variations in key adaptive traits, including floral and seed development, secondary metabolite biosynthesis, and plant responses to abiotic and biotic stresses. Indentification of gene family members in a newly sequenced genome is an essential step in functional and comparative genomics, enabling downstream analyses including gene expression profiling, selection pressure analyses, and evolutionary inferences.

Genome-wide gene family identification typically utilizes a combination of several established tools. Most gene family analysis studies rely on Hidden Markov Models (HMMs) profiling of specific domains to scan proteomes for candidate sequences, domain annotation tools to confirm the presence of conserved functional domains, and sequence similarity searches to validate candidates against curated references. Multiple sequence alignment, followed by phylogenetic reconstruction, provides an evolutionary framework for interpreting relationships among identified members. Although each of these steps is well-supported by established tools, integrating them requires manual handling of intermediate files, format conversions, output parsing, and consistent parameter management. These challenges can hinder reproducibility and scalability, particularly in collaborative projects or comparative analyses across multiple species.

There are dedicated pipelines that streamline the workflow of gene family analysis into a single reproducible framework. PlantTribes2 (Wafula et al. 2023) provides a scalable pipeline that classifies protein sequences into pre-computed orthologous clusters and supports downstream phylogenetic analyses. PangenePro (Fatima et al. 2025) is a recent shell-based tool that automates candidate identification, domain confirmation, and orthologous clustering, but it does not extend its functionality to phylogenetic inference. BAT (Steinbrenner 2024) supports flexible gene tree construction with functional annotation overlays with expression data and domain architecture. PlantTribes2 is automated using a modular Python/Perl script architecture, with primary delivery through Galaxy (Abueg et al. 2024). PangenePro is automated through a single shell script that calls a series of Python scripts sequentially. BAT’s automation is implemented via a Python package installed via pip. While each tool offers easy automation of its intended analytical scope, none provides execution-level infrastructure for step-level caching, parallel process execution, HPC scheduler integration, or fault-tolerant resumption, leaving reproducibility and portability dependent on the user’s computing environment. Workflow management systems such as Nextflow (Di Tommaso et al. 2017) address these limitations by providing a framework for defining, executing, and reproducing multi-step pipelines. Nextflow pipelines are portable, resumable on failure, and support multiple execution environments through containerization and package manager integration.

Here we present PhytoFam, a Nextflow DSL2 pipeline designed to address the above-mentioned limitations by automating the complete gene family identification workflow. PhytoFam integrates robust bioinformatics tools into a single reproducible framework, incorporating isoform-aware candidate filtering, multi-evidence sequence validation, and phylogenetic analysis with optional outgroup support. Although primarily designed to streamline gene family studies in plant proteomes, the pipeline is broadly applicable to any eukaryotic proteome using an HMM profile.

## 2 Materials and Methods

### 2.1 Software Dependencies

PhytoFam requires Nextflow v23.04 or later and Python v3.10 or later. Tool dependencies are managed through conda, Docker, or Singularity as described above. Specific versions used in development and testing are listed in Table 3.

### 2.2 Pipeline Overview

PhytoFam is implemented in Nextflow DSL2 and organized into six sequential analytical stages, each encapsulated as an independent module (Figure 1). The pipeline accepts a proteome in FASTA format, one or more HMM profile files, a reference protein sequence set of known gene family members, and optionally, outgroup sequences. All parameters are configurable *via* a centralized configuration file or command-line arguments, and the pipeline generates structured output directories for each analytical stage, along with a summary report.

**Figure 1.**
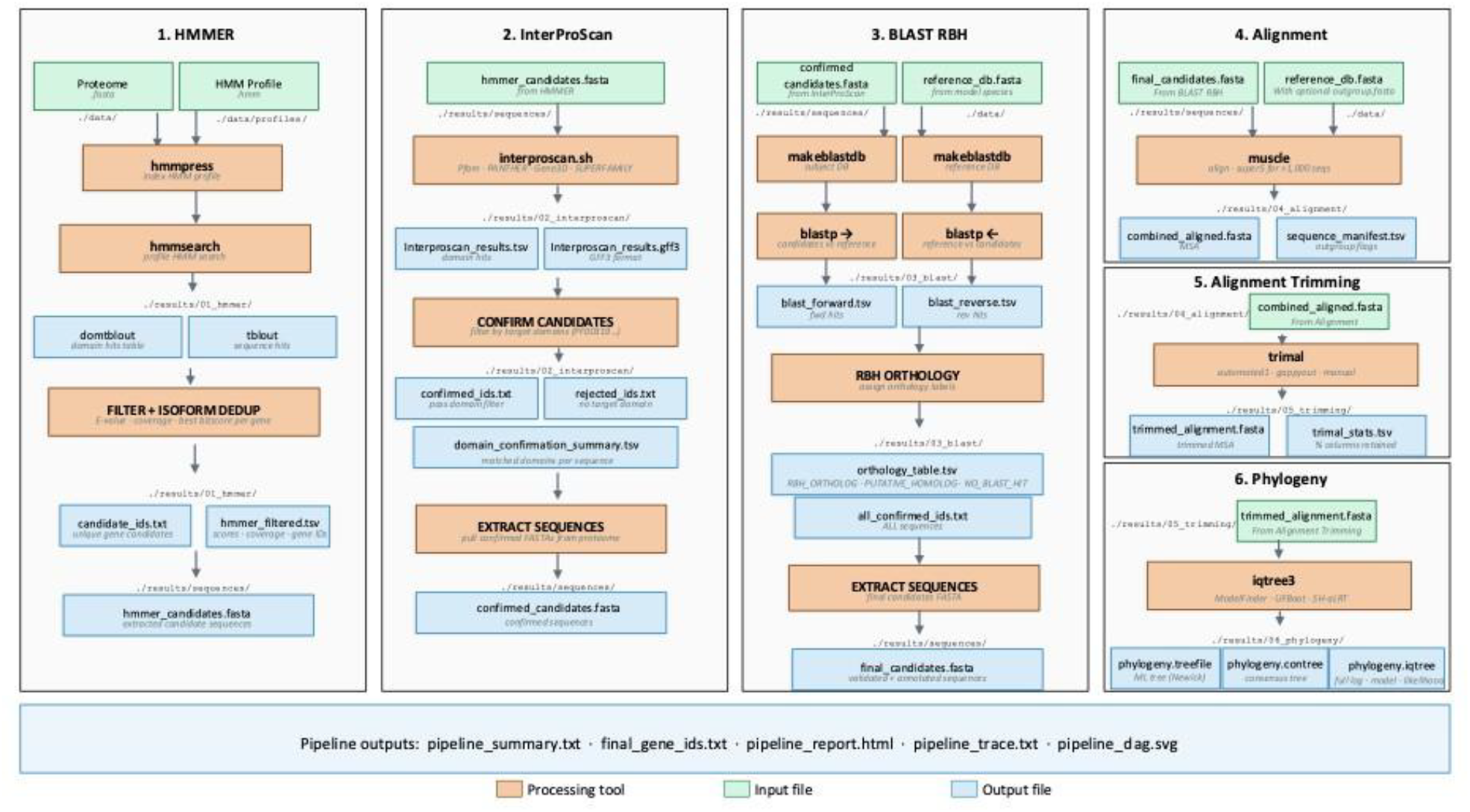
Schematic overview of the PhytoFam Nextflow pipeline for genome-wide identification of plant gene family. The pipeline comprises six modules executed sequentially: (1) HMMER hmmsearch scans the input proteome using user-supplied HMM profiles, followed by isoform deduplication retaining the highest-scoring representative per gene locus; (2) InterProScan validates candidate sequences based on the presence of user-specified Pfam domain accessions; (3) BLAST reciprocal best hit (RBH) analysis assigns orthology labels to all confirmed sequences; (4) MUSCLE aligns the candidate, reference, and (optional) outgroup sequences; (5) TrimAl trims the alignment by removing poorly aligned regions; and (6) IQ-TREE3 reconstructs maximum-likelihood phylogenetic tree with ultrafast bootstrap (UFBoot) and Shimodaira–Hasegawa approximate likelihood ratio test (SH-Alrt) support. Key input and output files, along with their directory paths, are shown. Color boxes depict input files (green), processing tools (orange), or output files (blue).

The six stages include: (1) HMMER-based candidate identification with isoform deduplication; (2) InterProScan domain confirmation; (3) BLAST reciprocal best hit (RBH) orthology assignment; (4) multiple sequence alignment with MUSCLE v5; (5) alignment trimming with TrimAl; and (6) maximum-likelihood phylogenetic inference with IQ-TREE3. Each stage is described in the following sections.

#### Stage 1: HMMER-Based Candidate Identification and Isoform Deduplication

The first stage uses HMMER v3.4 (Potter et al. 2018) to search the target proteome against one or more user-supplied HMM profiles using hmmsearch. If multiple profile files are provided, they are concatenated and indexed into a database prior to searching. Candidate sequences are retained if their full-sequence E-value and domain-level HMM coverage satisfy user-defined thresholds (default: E-value ≤ 1 × 10^−5^; HMM coverage ≥ 0.5). These thresholds ensure both statistical confidence and sufficient alignment to conserved model features of the HMM profile.

To mitigate redundancy introduced by alternative splicing, which commonly results in multiple predicted protein isoforms per gene locus, PhytoFam incorporates an automated isoform deduplication procedure. Gene identifiers are inferred from sequence IDs using a rule-based approach that strips common isoform suffixes (e.g., .1, .2, .1.1, -01) present in genome annotation conventions. For each inferred gene locus, the isoform with the highest bitscore is retained, with ties resolved by E-value (ascending) and then HMM profile coverage (descending), thereby favoring sequences with stronger statistical support and more complete domain representation. The selected representative and its corresponding gene identifier are recorded in the output file *hmmer_filtered*.*tsv*.

#### Stage 2: InterProScan Domain Confirmation

Candidate sequences identified by HMMER are evaluated for domain confirmation using InterProScan v5.69.101 (Jones et al. 2014) . InterProScan is run against multiple member databases, including Pfam, PANTHER, Gene3D, SUPERFAMILY, PRINTS, and ProSite profiles, with Gene Ontology term and pathway annotation enabled to provide functional context for detected features. Candidates are confirmed if at least one hit is detected to a user-specified set of Pfam or InterPro accessions (*--target_domains)*, provided as a comma-separated list.

Confirmed and rejected sequence IDs are written to separate output files. The confirmation summary table records the matched domain accessions for each sequence, providing an audit trail for downstream analyses and debugging.

#### Stage 3: BLAST Reciprocal Best Hit Orthology Assignment

Following domain confirmation, all confirmed candidate sequences are annotated with orthology information using a BLAST reciprocal best hit (RBH) approach with NCBI-BLAST (version 2.17.0) (Altschul et al. 1990; Camacho et al. 2009) . All sequences confirmed by InterProScan proceed to alignment and phylogenetic analysis regardless of their RBH BLAST result. This is because lineage-specific gene family members or highly diverged sequences may lack a clear RBH to the reference set yet remain valid gene family members.

The RBH analysis proceeds in two directions: candidate sequences are searched against the reference protein database (forward BLAST), and reference sequences are searched against the full target proteome (reverse BLAST), both using blastp with a consistent E-value threshold and minimum identity and query coverage filters. The orthology assignment for each confirmed sequence is classified into one of three categories: RBH_ORTHOLOG (strict reciprocal best hit), PUTATIVE_HOMOLOG (significant forward hit without reciprocation), or NO_BLAST_HIT (no significant hit at the defined thresholds). These annotations are recorded in orthology_table.tsv.

#### Stage 4: Multiple Sequence Alignment

Confirmed candidate sequences are merged with the reference sequence set and, if provided, user-specified outgroup sequences, prior to multiple sequence alignment. Sequence merging is performed with source tracking: each sequence is labeled as a candidate, reference, or outgroup in the output sequence manifest (sequence_manifest.tsv), enabling downstream identification of outgroup taxa for tree rooting. Duplicate sequence identifiers are resolved by retaining the first occurrence in order of priority (candidates > references > outgroup).

Alignment is performed using MUSCLE (version v5.3) (Edgar 2021). For datasets exceeding 1,000 sequences, the pipeline automatically switches from the standard -align algorithm to the super5 algorithm, which is optimized for large-scale alignments. The algorithm selection is logged in muscle.log for reproducibility.

#### Stage 5: Alignment Trimming

The multiple sequence alignment is trimmed using TrimAl (version v1.5.1) (Capella-Gutiérrez et al. 2009) to remove poorly aligned or gap-rich columns that could introduce noise into phylogenetic inference. The trimming strategy is user-configurable *via* the *--trimal_method* parameter, with supported options including *automated1* (default), *gappyout, strict, strictplus*, and *manual*. With the manual method, users can specify the additional -gt (*--trimal_gt*) argument as needed.

Pre- and post-trimming alignment statistics, including the number of columns retained and the percentage of alignment length preserved, are recorded in trimal_stats.tsv. An HTML visualization of the trimmed alignment is also produced for manual inspection.

#### Stage 6: Maximum-Likelihood Phylogenetic Inference

Maximum-likelihood phylogenetic inference is performed using IQ-TREE3 (version v3.1.2) (Wong et al. 2026). By default, the substitution model is selected automatically using ModelFinder (Kalyaanamoorthy et al. 2017) (*-m TEST*), which evaluates a range of protein substitution models and selects the best-fit model based on the Bayesian Information Criterion (BIC). Users may alternatively specify a model explicitly *via --iqtree_model*. Branch support is assessed using ultrafast bootstrap (UFBoot) with 1,000 replicates by default, supplemented by SH-aLRT with 1,000 replicates, providing two complementary measures of clade support. IQTREE3 is run with five independent tree searches (*--runs 5*) by default to reduce the risk of entrapment in local optima.

The pipeline produces an unrooted maximum-likelihood tree; it is up to the user to root it as desired. When outgroup sequences have been included in the alignment, the sequence_manifest.tsv file produced in Stage 4 provides the outgroup sequence identifiers needed to root the tree in downstream visualization tools such as FigTree (Rambaut 2009), Interactive Tree of Life (iTOL) (Wong et al. 2026), ggtree (R) (Yu et al. 2017), or ETE3 (Python) (Huerta-Cepas et al. 2016).

### 2.3 Implementation

#### 2.3.1 Pipeline Architecture and Nextflow Configuration

PhytoFam is implemented using Nextflow DSL2, in which each analytical stage is defined as an independent module in a dedicated .*nf* file. This modular architecture allows individual processes to be updated, replaced, or reused independently of the broader workflow. The entry point for the pipeline is *main*.*nf*, which imports all modules and defines the workflow channel logic. All parameters, resource allocations, and execution profiles are centralized in *nextflow*.*config*.

##### Parameter Configuration

The *nextflow*.*config* file defines all user-facing parameters in a params block. Required parameters include the paths to the target proteome (*--proteome*), HMM profiles (*-hmm_profiles)*, reference sequences (*--reference_db*), and the target domain Pfam accessions for InterProScan confirmation (*--target_domains*). Optional parameters include the outgroup sequence file (*--outgroup*), and all tool-specific thresholds. Default values are provided for all optional parameters and are listed in Table 2.

**Table 1.** Summary of software dependencies and versions.

| Software | Version Tested | Reference |
| --- | --- | --- |
| Nextflow | 25.04.2 | Di Tommaso et al. (2017) |
| HMMER | 3.4 | Potter et al. (2018) |
| InterProScan | 5.59_91.0 | Jones et al. (2014) |
| BLAST+ | 2.17.0 | Altschul et al. (1990); Camacho et al. (2009) |
| MUSCLE | 5.3 | Edgar (2021) |
| TrimAl | 1.5.1 | Capella-Gutiérrez et al. (2009) |
| IQ-TREE3 | 3.1.2 | Wong et al. (2026) |

**Table 2.**
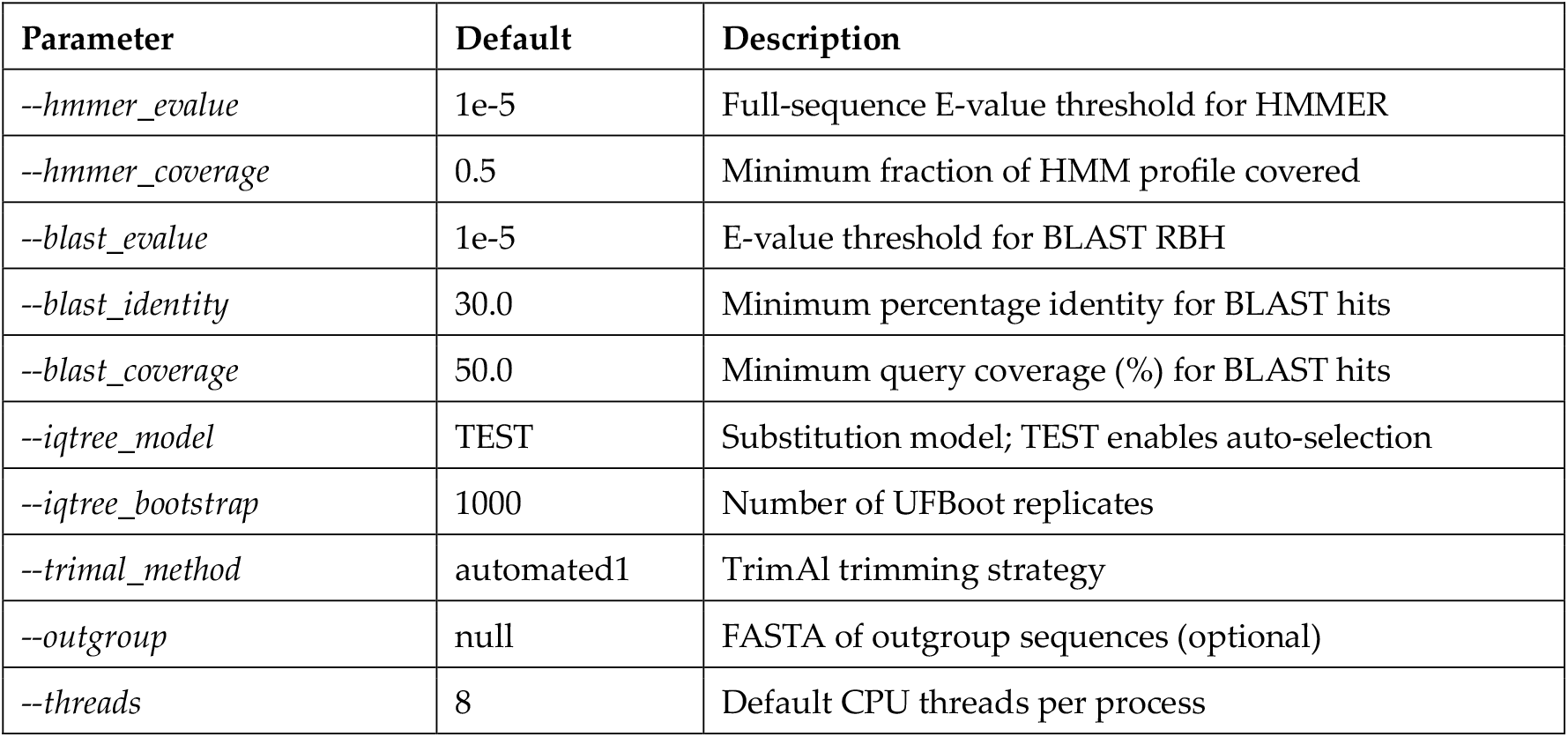
List of PhytoFam configurable parameters and default values.

**Table 3.** List of primary output files produced by PhytoFam.

| File | Description |
| --- | --- |
| 01_hmmer/hmmer_filtered.tsv | HMMER hits passing filters; includes selected isoform and gene ID per locus |
| 01_hmmer/candidate_ids.txt | Sequence IDs of deduplicated HMMER candidates |
| 02_interproscan/domain_confirmation_summary.tsv | Per-sequence domain confirmation status and matched accessions |
| 03_blast/orthology_table.tsv | RBH orthology classification for all confirmed sequences |
| 04_alignment/sequence_manifest.tsv | Source label (candidate/reference/outgroup) and outgroup flag per sequence |
| 05_trimming/trimal_stats.tsv | Alignment column statistics before and after trimming |
| 06_phylogeny/phylogeny.treefile | Maximum-likelihood tree in Newick format |
| pipeline_summary.txt | Candidate counts at each filtering stage; links to key output files |
| final_gene_ids.txt | Final list of confirmed gene family member IDs |

##### Resource Management

Process-level resource allocation in PhytoFam is managed through process labels defined in *nextflow*.*config*. Four labels are defined with increasing resource tiers: low_cpu (2 CPUs, 4 GB RAM), medium_cpu (user-defined threads, 16 GB RAM), high_cpu (user-defined threads, 32 GB RAM), and high_mem (user-defined threads, 64 GB RAM). Each process is assigned an appropriate label based on its expected computational requirements. HMMER, BLAST, MUSCLE, and IQ-TREE are assigned medium_cpu or high_cpu labels due to their support for parallelization, while lightweight parsing and filtering processes are assigned low_cpu. Processes are configured to retry up to two times for exit codes associated with memory or compute node failures (exit codes 143, 137, 104, 134, 139). Users are welcome to make necessary changes in the config file if their computational infrastructure uses different partition names or memory configurations.

##### Execution Profiles

*PhytoFam* provides pre-configured execution profiles for common deployment scenarios. The conda profile enables automatic environment creation using Mamba (with fallback to conda) (Anaconda 2016; Grüning et al. 2018), with tool-specific environments defined per process group. The Docker (Merkel 2014) and Singularity (Kurtzer et al. 2017) profiles enable containerized execution using Biocontainer (Leprevost et al. 2017) images. To note,

InterProScan requires a local installation prior to running the current PhytoFam version. Users can download and install InterProScan from the official EBI repository (https://github.com/ebipf-team/interproscan) and specify the path to the interproscan.sh executable via the *-interproscan_bin* parameter or ensure it is available on the system *$PATH* or load the module in the environment (*module load interproscan*) prior to running the pipeline. All other dependencies are managed automatically through the pipeline’s conda, Docker, or Singularity profiles.

The Slurm and LSF profiles configure submission to HPC schedulers with configurable queue and account settings. Profiles may be combined at runtime (e.g., *-profile slurm,conda*) to enable scheduler-aware execution with conda-managed dependencies.

#### 2.3.2 Output Structure

All pipeline outputs are written to a user-specified directory (default: results/), organized by analytical stage. Table 3 summarizes the primary output files produced by PhytoFam.

#### 2.3.3 Case Study: Genome-Wide Identification of MADS Box Gene Family in White Mulberry

The white mulberry (*Morus alba* cv. Heyebai) genome (GenBank Assembly GCA_012066045.3) retrieved from NCBI was used to survey members of the MADS Box gene family. The genome was annotated with the BRAKER3 pipeline (Gabriel et al. 2024). BRAKER3 uses ProtHint for protein-genome alignments, GeneMark-EP+ (Brůna et al. 2020) for evidence-supported gene prediction, and AUGUSTUS (Stanke et al. 2006) for final gene model training and refinement. After confirming the assembly was soft-masked, BRAKER was executed in protein-guided mode (BRAKER2) using the Viridiplantae.fa protein dataset from OrthoDB (Kuznetsov et al. 2023) as external evidence, with the *--prot_seq* option and *--softmasking* enabled. The workflow was executed inside a Singularity container (braker3.sif). The annotated proteome (Supplementary File 1) had the characteristic AUGUSTUS-type gene IDs (example: g1234.t1, g1234.t2), where the characters before the period (.) denote the gene ID and the ones after the period denote the isoform corresponding to the gene ID. The proteome had 67,792 protein sequences and was used as the target proteome (proteome) in the workflow. 105 *Arabidopsis thaliana* MADS Box protein sequences retrieved from The Arabidopsis Information Resource (TAIR) (Supplementary File 2) (Reiser et al. 2024) were used as the reference (--reference_db). HMM profile of the SRF-type transcription factor (DNA-binding and dimerisation domain) (PF00319.hmm) was retrieved from the InterPro database (Blum et al. 2025) (retrieved May 18, 2026) and was used as the HMM profile (--hmm_profiles), and the target domain argument (--target_domains) was defined as PF00319. Considering the diversity observed in plant MADS-box protein sequences (clade and lineage-specific), the TrimAl trimming parameter was set to a more flexible -gt 0.2 (*-trimal_method manual --trimal_gt 0*.*2*), i.e., a column is retained if at least 20% of sequences have a residue there. The pipeline was run without an outgroup using the conda profile (*-profile conda*) in an interactive (*srun*) session on a single bigmem node (Dell PowerEdge R750, Intel Xeon Gold 6342 CPU @ 2.80 GHz) with 40 allocated CPU cores and 2TB RAM, with individual processes allocated 2-8 threads as defined by the pipeline resource configuration. All analyses were performed on the Innovator High Performance Computing cluster at South Dakota State University, running Rocky Linux 9. The command used to run the pipeline was:

*nextflow run* .*/PhytoFam/main*.*nf -profile conda --proteome data/morus_proteome*.*fasta -hmm_profiles data/profiles/PF00319*.*hmm --reference_db data/arabidopsis_MADS*.*fasta -target_domains PF00319 --trimal_method manual --trimal_gt 0*.*2 --outdir results -resume*

## 3 Results

### 3.1 Computational Summary

The complete pipeline execution time of analyzing the MADS Box gene family in the *M. alba* proteome was 1 hour 10 minutes and 8 seconds, which corresponded to 9 CPU hours (h). The .txt file with the pipeline trace data is provided as Supplementary File 3. In terms of runtime (real273 time), IQTREE3 was the lengthiest process (1 h), followed by MUSCLE_ALIGN (3.6 minutes) and INTERPROSCSAN (1.4 minutes). INTERPROSCAN was the hungriest stage in the entire pipeline, using 4.5 gigabytes (GB) (Resident Set Size, RSS) of peak memory, followed by MUSCLE_ALIGN at 447 megabytes (MB) and IQTREE3 at 336 MB. The ‘Phylogenetics’ 277 portion of the pipeline used most of the 9 CPU hours used (8.30 h), followed by the ‘Alignment’ stage (0.48 h) and ‘InterProScan’ (0.19 h) (Figure 2).

**Figure 2.**
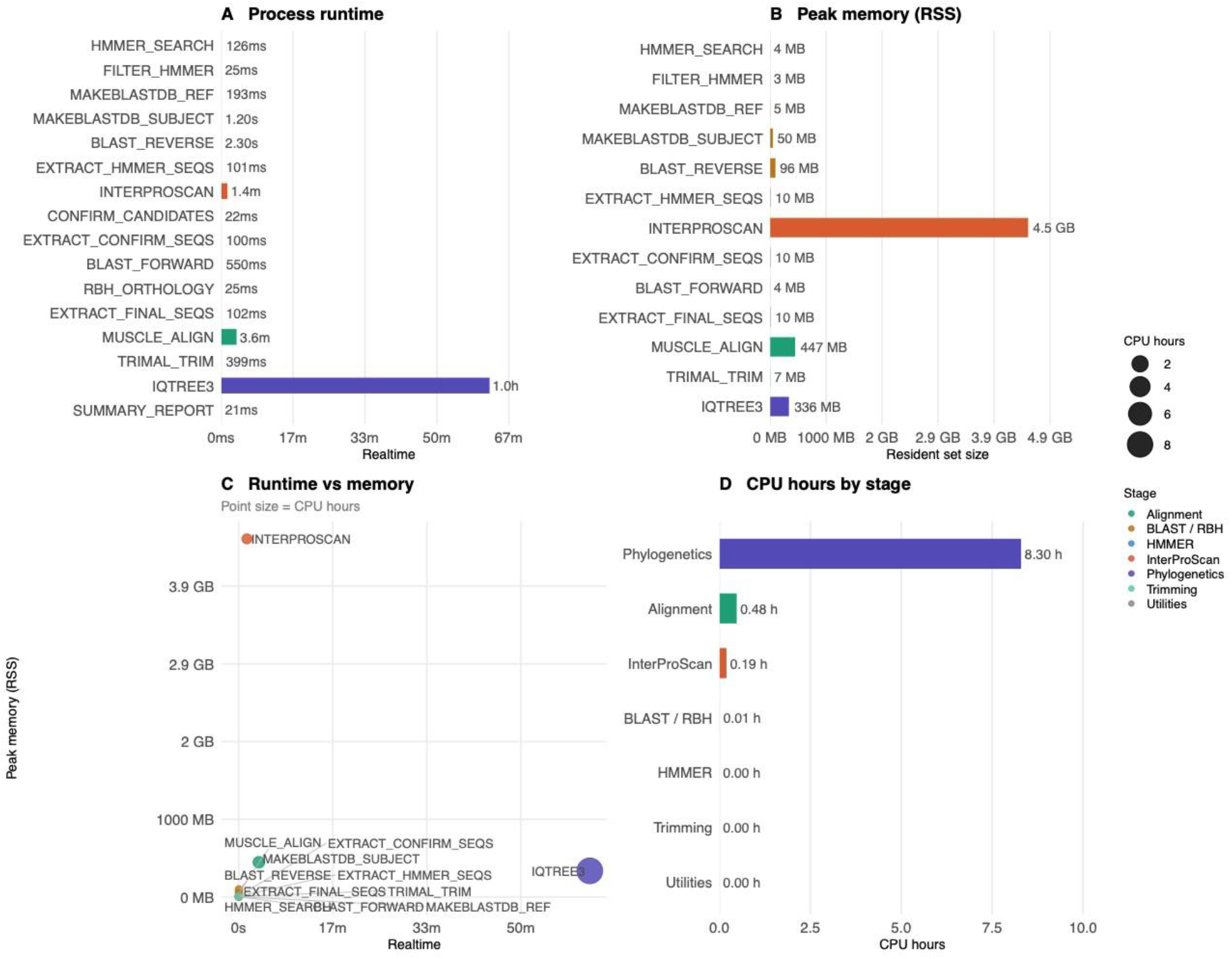
Runtime and resource usage of the PhytoFam pipeline assessed on the *Morus alba* proteome for MADS-box gene family identification. (A) Wall clock/real-time for each of the 16 pipeline processes. IQ-TREE3 dominated the total runtime at approximately 1 hour, accounting for 88% of the wall time. (B) Peak resident set size (RSS) for each process. InterProScan exhibited the highest memory footprint (4.5 GB), attributable to the in-memory loading of domain reference databases, while IQ-TREE3 and MUSCLE consumed 336 MB and 447 MB, respectively. (C) Scatter plot of real-time versus peak memory per process, with point size scaled to CPU hours consumed, illustrating the distinction between CPU-bound (IQ-TREE3) and memory-bound (InterProScan) processes. (D) Total CPU hours per pipeline stage. Phylogenetic inference accounted for 8.30 of the 9h total CPU hours consumed. All analyses were run on a single high-memory node (Dell PowerEdge R750, Intel Xeon Gold 6342 @ 2.80 GHz, 40 allocated cores) on the Innovator HPC cluster at South Dakota State University (Rocky Linux 9).

### 3.2 Gene Family Analysis Report

In the first stage, HMMER was executed with default thresholds of E-value ≤ 1×10^−5^ and HMM coverage ≥ 0.5, yielding 75 raw hits. The pipeline’s isoform deduplication step identified and removed eight redundant sequences representing alternative isoforms of the same gene loci, retaining 67 unique gene candidates/hits for downstream analysis. These hits were then passed to InterProScan, which confirmed the presence of the MADS-box domain in all of them. The pipeline then performed BLAST-based orthology assignment, classifying 21 candidates as orthologs through reciprocal best hits with *A. thaliana* MADS-box proteins, 36 as putative homologs based on significant forward hits only, and the remaining 10 as having no significant BLAST hit against the reference set, suggesting they may represent lineage-specific or diverged members.

The *M. alba* hits, and 105 *A. thaliana* MADS-box proteins were merged and aligned using MUSCLE v5. TrimAl trimmed the alignment using a gap threshold of 0.2, retaining only 20% of alignment columns. This low retention rate is likely due to the structural divergence between Type I and Type II MADS-box proteins, where Type I proteins possess only a divergent MADS domain while Type II (MIKC-type) proteins additionally contain the I, K, and C domains (De Bodt et al. 2003), producing extensive gap-rich regions when both MADS domain types are aligned together. The trimmed alignment was passed to IQ-TREE3, where ModelFinder evaluated 224 substitution models and selected Q.INSECT+F+G4 as the best-fit model based on the BIC. Among five independent tree searches, Run 5 produced the best log-likelihood score of −64,276.980. The resulting tree file was rooted at the midpoint, annotated, and visualized in iTOL v6 (Figure 3). Phylogenetic classification of the *M. alba* hits revealed 40 MIKC, 13 Malpha, five M-beta, five M-delta, and four M-gamma-type members. These numbers are consistent with those previously reported for the *Morus notabilis* genome: 33 MIKC, 11 Malpha, two M-beta, four M-delta, and four M-gamma (Luo et al. 2018).

**Figure 3.**
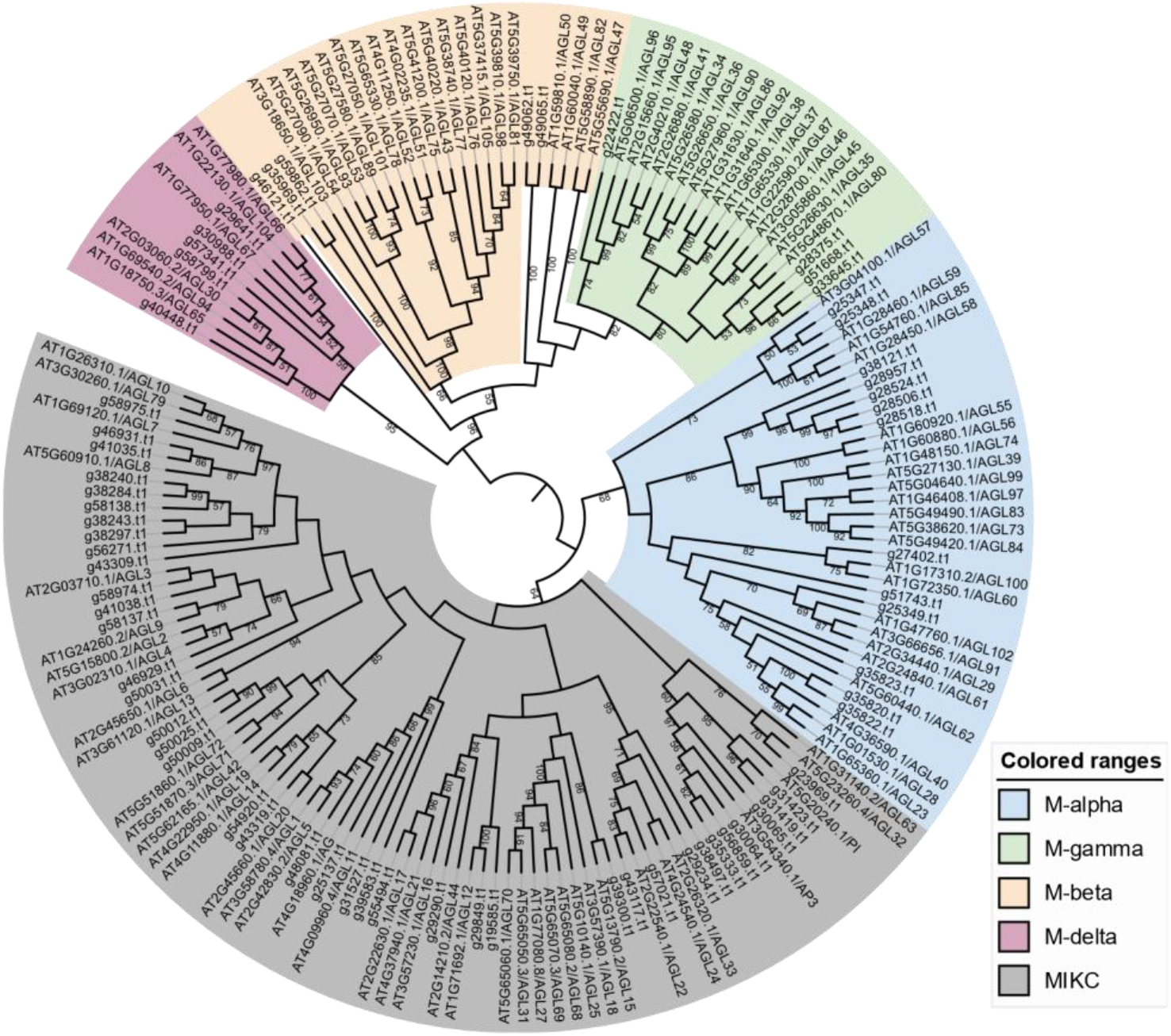
Maximam - likelihood phylogenetic tree of MADS-box proteins identified in *MORUS alba* and reference proteins of *Arabidopsis thaliana*. The tree was generated using IQ-TREE3 with the Q.INSECT+F+G4 substitution model selected by ModelFinder based on the Bayesian Information Criterion. Branch support values represent ultrafast bootstrap (UFBoot) from 1,000 replicates. The tree was rooted at the midpoint and visualized in iTOL v6. Colored ranges indicate subfamily classifications: M-alpha (blue), M-gamma (green), M-beta (peach), M-delta (pink), and MIKC-type (grey). *A. thaliana* sequences are prefixed with AT; *M. alba* sequences are prefixed with g.

## 4 Discussion

PhytoFam provides a reproducible, portable workflow for genome-wide identification of plant gene families. This workflow integrates multiple lines of evidence and covers the complete analytical process from proteome to phylogenetic tree. By combining HMM-based detection, domain-level confirmation, and sequence similarity-based orthology assignment, PhytoFam reduces false positives that can plague downstream analyses.

The pipeline is designed to be a practical automated tool for plant genomics research, but we expect users to evaluate the outputs of each stage critically, including E-values, domain-selection sensitivity, and alignment-trimming parameters, depending on the gene families and proteomes under investigation. Users are expected to supply appropriate HMM profiles and target domain accessions for the gene family of interest, and to interpret the resulting phylogenetic tree using domain knowledge of the family. It can be useful to work with the different parameters (Table 2) for a more exhaustive initial search, especially when the candidates in the initial search have Evalues and coverage values borderline to the threshold, to ensure that divergent members are not omitted from downstream analyses. Furthermore, not all domains/gene families of interest have a Pfam domain accession, and an HMM profile hosted by the InterPro database. In such cases, users are encouraged to build their own HMM profiles using the HMMER tool. In these cases, users are also encouraged to set their --target_domains flag to the domain’s accession ID in databases other than Pfam. By default, the memory-intensive step of InterProScan has been programmed to run with a minimum of 32 GB of memory for a faster pipeline completion, and the pipeline will stop if the available memory is lower than that number. Users are encouraged to edit that parameter in the *nextflow*.*config* file if they are limited by computational resources.

There are pipelines for gene family identification and analysis, such as PangenePro (Fatima et al. 2025) and BAT (Steinbrenner 2024). PangenePro uses the BLAST algorithm to identify gene family members across multiple genomes/ecotypes of the same species. PhytoFam starts off with an HMM-based ortholog identification, which has better sensitivity to distant orthologs than BLAST (Potter et al. 2018). PangenePro does not have an in-built phylogenetic inference step, unlike PhytoFam. PlantTribes2 (Wafula et al. 2023) is another tool for genome-wide gene family analyses like PhytoFam, but both have different philosophies. PlantTribes2 is scaffold-based and groups BLAST and HMMER-based candidates into pre-computed orthogroups. PhytoFam does not have an orthogroup dependency, which makes it more suitable for new genomes of nonmodel species. PhytoFam has InterProScan-based filtering built in for each candidate, whereas PlantTribes2 has InterPro/Pfam annotations for each orthogroup, not for individual candidates. PlantTribes2 has an edge over PhytoFam in its deployment via the Galaxy Suite (Abueg et al. 2024), making it more accessible to users without strong informatics experience. PlantTribes2 also has a built-in genome duplication inference tool with genome-scale evolutionary analysis, which is beyond the scope of PhytoFam.

Future development directions include support for genome-level GFF3 coordinate extraction for identified members, multi-species comparative analysis modes, integration with synteny analysis tools, and analysis of conserved motifs. The inclusion of a step to build the HMM profile from a user-supplied set of reference sequences will be part of future releases.

## Acknowledgement

PhytoFam uses code and infrastructure developed and maintained by the nf-core community. The authors thank the Research Cyberinfrastructure team at South Dakota State University for providing High-Performance Computing (HPC) services required for this study. Pipeline scripts were debugged and formatted with assistance from Claude (Anthropic).

## Author Contributions

Sanam Parajuli (Conceptualization, Investigation, Visualization, Methodology, Formal Analysis, Writing – original draft, Writing –review and editing), Bibek Adhikari (Writing –review and editing), Anne Fennell (Writing –writing, review and editing), Madhav P. Nepal (Conceptualization, Investigation, Funding acquisition [lead], Supervision [lead], Writing –review and editing)

## Supplementary Data

(available at https://doi.org/10.6084/m9.figshare.33071060)

Supplementary File 1 *Morus alba* proteome FASTA file

Supplementary File 2 *Arabidopsis thaliana* MADS-box proteins FASTA file

Supplementary File 3 Pipeline Trace txt file

## Availability

PhytoFam is released under the GNU General Public License v3.0. Source code and documentation are available at https://github.com/sanamparajuli/PhytoFam. The pipeline can be installed locally with the git clone mechanism or executed without local installation with the nextflow pull mechanism.

